# MSdb: an integrated expression atlas of human musculoskeletal system

**DOI:** 10.1101/2022.11.23.517756

**Authors:** Junxin Lin, Ruonan Tian, Ziwei Xue, Dengfeng Ruan, Pengwei Chen, Yiwen Xu, Chao Dai, Weiliang Shen, Hongwei Ouyang, Wanlu Liu

**Affiliations:** Department of Orthopedic of the Second Affiliated Hospital of Zhejiang University School of Medicine, Zhejiang University, Hangzhou, Zhejiang, China; Zhejiang University-University of Edinburgh Institute, Zhejiang University School of Medicine, Hangzhou, China; Dr. Li Dak Sum & Yip Yio Chin Center for Stem Cells and Regenerative Medicine, Zhejiang University School of Medicine, Hangzhou, China; Key Laboratory of Tissue Engineering and Regenerative Medicine of Zhejiang Province, Hangzhou, Zhejiang, China; Department of Sports Medicine, Zhejiang University School of Medicine, Hangzhou, China; China Orthopedic Regenerative Medicine Group (CORMed), Hangzhou, China; Alibaba-Zhejiang University Joint Research Center of Future Digital Healthcare, Zhejiang University, Hangzhou

## Abstract

We introduce MSdb (https://www.msdb.org.cn), a database for visualization and integrated analysis of next-generation sequencing data from human musculoskeletal system, along with manually curated patient phenotype data. Systematic categorizing, standardized processing and freely accessible knowledge enables the reuse of public data. MSdb provides various types of analysis, including sample-level browsing of metadata information, gene/miRNA expression, and single-cell RNA-seq dataset. In addition, MSdb also allows integrated analysis for cross-samples and cross-omics analysis, including customized differentially-expressed gene/microRNA analysis, microRNA-gene network, scRNA-seq cross-sample/disease integration, and gene regulatory network analysis.

## Main

The prevalence and burden of musculoskeletal (MSK) disorders are extremely high all over the world^1,2^. With the advancement of next-generation sequencing (NGS), a huge amount of sequencing data has been generated, which accelerated the research of pathological mechanisms and the development of novel therapeutic approaches for MSK disorders^3-6^. However, dispersed distribution of relevant data sets among different repositories makes it challenging to analyze and compare them in a uniform way. Therefore, we developed the human musculoskeletal system database (MSdb), an integrated expression atlas specifically for the human MSK system, containing 33 diseases, 126 projects, 3,398 transcriptomes and microRNAomes at bulk level, as well as 2,833,779 transcriptomes at single-cell level (Fig. 1a and Supplementary Fig. 1). MSdb incorporates cross-repository metadata into controlled vocabulary and uniform format, enabling efficient retrieval of sample information (Supplementary Table). MSdb provides multiple built-in data exploration and analysis functionalities, including gene/microRNA expression browsing, customized differentially-expressed genes/miRNAs analysis, integrated microRNA-gene interaction networks, as well as integrated single-cell expression atlas and cell type-specific gene regulatory networks analysis (Supplementary Fig. 2 and Supplementary Video). Furthermore, MSdb allows downloading of processed data sets and publication-quality plots, offering wet lab scientists powerful tools to browse and re-analyze the public datasets without technical barriers.

**Fig. 1.**
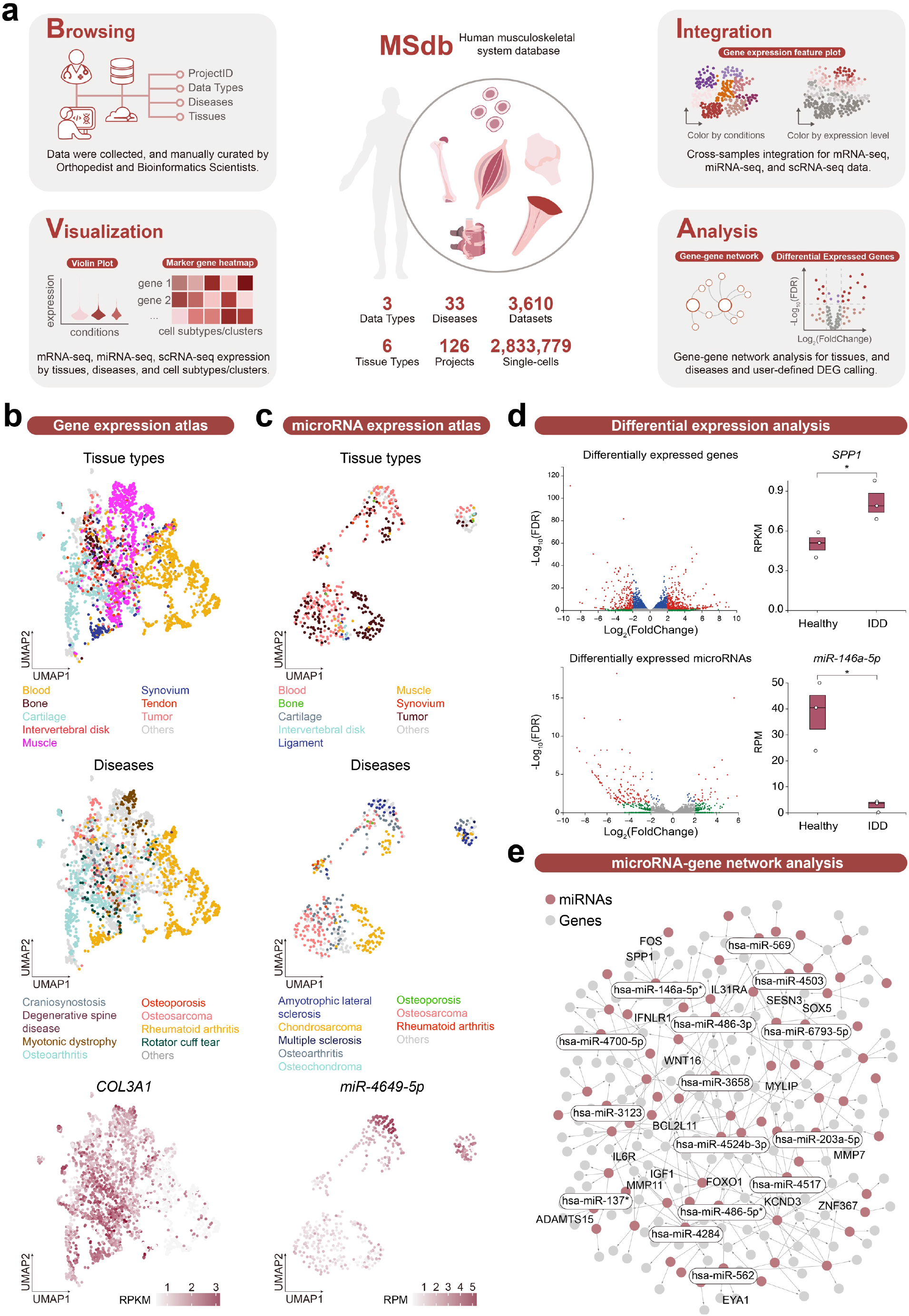
MSdb framework and illustrative data analysis. **a**, Overview of MSdb. MSdb is a comprehensive database of next-generation sequencing data on human musculoskeletal system tissues and cells, enhanced with manually curated patient phenotypes, advanced analysis and visualization tools. **b, c**, UMAP plots showing gene (**b**) and microRNA (**c**) expression atlas in MSdb. All gene or microRNA expression data in MSdb were used for clustering, respectively. Samples are colored by tissue types (top), diseases (middle) and *COL3A1*/*miR-4649-5p* expression levels (bottom). **d**, Volcano plots and box plots showing dysregulated genes (top) or microRNAs (bottom) between healthy (n=3) and degenerative (n=3) intervertebral disks. RPKM: reads per kilobase per million mapped reads; RPM: reads per million mapped reads. *: FDR < 0.01. **e**, microRNA-gene interaction network built with down-regulated microRNAs and up-regulated genes in degenerative intervertebral disks. Red dots represent the microRNAs and grey dots represent the genes. Complete and interactive plots of the network are available online on MSdb database.

MSdb enables users to retrieve sample information via consistent and validated metadata curated by orthopedists and bioinformatics scientists (Supplementary Table). Users can search the database using multiple parameters like project identifier, diseases, and tissue types in order to find data sets that match their interests (Supplementary Fig. 3a). In metadata, four types of information are available: (i) data set and publication identifiers, (ii) patient phenotypes, (iii) sample information and (iv) data pre-processing summary. These items enable users to evaluate the biological meaning, clinical relevance and data quality of the samples. Summary statistics of the metadata are also presented to show global patterns of the studies (Supplementary Fig. 4).

At bulk level, MSdb integrates information at two aspects: (i) cross-tissue integration, (ii) cross-omics integration. For cross-tissue integration, samples were initially integrated by projects to generate gene or microRNA expression atlas, and then labelled by their tissue types, diseases, cell types, and tissue positions. Users may explore the expression of genes or microRNAs by these labels (Supplementary Fig. 5 and Supplementary Fig. 6). In Fig. 1b, c, uniform manifold approximation and projection (UMAP) plots show the integrated expression atlas of bulk RNA-seq or microRNA-seq data in MSdb, and samples are colored by representative tissue types and diseases. Feature plots and violin plots show that *COL3A1*, a connective tissue marker, was pervasively expressed in MSK tissues as expected (Fig. 1b and Supplementary Fig.5). Cerebrospinal fluid from amyotrophic lateral sclerosis (ALS) patients showed enrichment of the potential diagnostic marker miR-4649-5p (Fig. 1c and Supplementary Fig.6)^7^. For cross-omics integration, MSdb enables users to analyze bulk RNA-seq and microRNA-seq data side-by-side and integrate the results to predict disease-related gene expression regulatory mechanisms. MSdb’s differential expression analysis module allows users to choose two groups of samples for comparison (Supplementary Fig. 3b). An interactive volcano plot can be used to explore differentially expressed genes or microRNAs between user-defined groups, and the expression level of a specific gene or microRNA can be queried and will be displayed in the box plot (Supplementary Fig. 3b). By integrating differentially expressed genes and microRNAs between normal and pathological intervertebral disks, we constructed a microRNA-gene interaction network (Fig. 1d, e). The network revealed previous reported (indicated by asterisk) and unanticipated disease-associated microRNAs and their potential gene targets, which can be interactively explored on our website (Fig. 1e). We demonstrated that the expression level of *miR-146a-5p* was down-regulated, while its target SPP1 was up-regulated in degenerative intervertebral disks when compared to healthy control (Fig. 1d, e). Since abnormal expression of SPP1 plays a critical role in the pathological process of intervertebral disk degeneration (IDD)^8^, our analysis indicated that targeting SPP1 with *miR-146a-5p* mimic might be a therapeutic method to counteract disk degeneration. Collectively, MSdb offers users with integrated expression atlas and useful data analysis functionalities to understand gene function and regulation in homeostasis and diseases.

In recent years, the emergence of single-cell technology permits researchers to discover new and possibly unexpected biological findings relative to bulk-level profiling. Intriguingly, MSdb contains a wealth of single-cell RNA sequencing (scRNA-seq) data and provides a suit of functionalities for users to explore gene expression at single-cell level. For each scRNA-seq dataset, we provide textual and graphical representations for sample information, quality control metrics, unsupervised cell clustering, reference-based cell subtype prediction, as well as marker genes for each cell type (Supplementary Fig. 7). In a single-cell profiling of synovial tissue from a female patient with rheumatoid arthritis, major cell types, such as fibroblasts, macrophages, T cells and monocytes were identified and the specific *CD68* expression in the predicted macrophages (cluster 2) indicated the reliability of cell clustering and automated cell type prediction (Supplementary Fig. 7a-h). For more sophisticated cell type clustering and annotation, users can adjust leiden resolution for cell clustering and change reference dataset for cell type prediction (Supplementary Fig. 7i). To help with manual annotation, a heatmap and a table of marker genes are displayed and available for download. (Supplementary Fig. 7j, k).

We also implemented an in-house variational autoencoder (VAE) based deep-learning framework to facilitate the integrative analysis for scRNA-seq data sets from different studies. Fig. 2 displays the result of integrated analysis of single-cell gene expression data from healthy, osteoarthritis (OA), rheumatic arthritis (RA) and undifferentiated arthritis (UA) patients. Using the VAE model, we were able to remove batch differences and integrate heterogeneous data from different studies (Fig. 2b). The cell types could be identified by the known marker genes, whose expression patterns support that our integration method appropriately aligned gene expression for each cell types (Fig. 2a, c). Interestingly, we observed a distinct distribution pattern of fibroblasts from OA and RA patients (Fig. 2b). Differential expression analysis revealed that *IGFBP3* and *LOX* were more enriched in OA-derived fibroblast when compared to RA-derived fibroblast (Fig. 2d). *IGFBP3* and *LOX* were involved in extracellular matrix remodeling, which may determine synovial fibrosis and may be associated with the clinical symptoms of pain, hyperalgesia, and stiffness in osteoarthritis^9^. It was also noted that *CD74* and *HLA-DRA* were specifically expressed in RA fibroblasts, demonstrating an inflammatory state of fibroblasts that have been reported to be a major source of pro-inflammatory cytokines and highlighted as a potential therapeutic target in RA (Fig. 2d)^10,11^. Moreover, we inferred gene regulatory networks (GRNs) using a previously published deep regenerative model for each cell types^12^. Users may interactively visualize the GRNs in our database to explore the complicated molecular interactions governing potential cell identity (Supplementary Fig. 8).

**Fig. 2.**
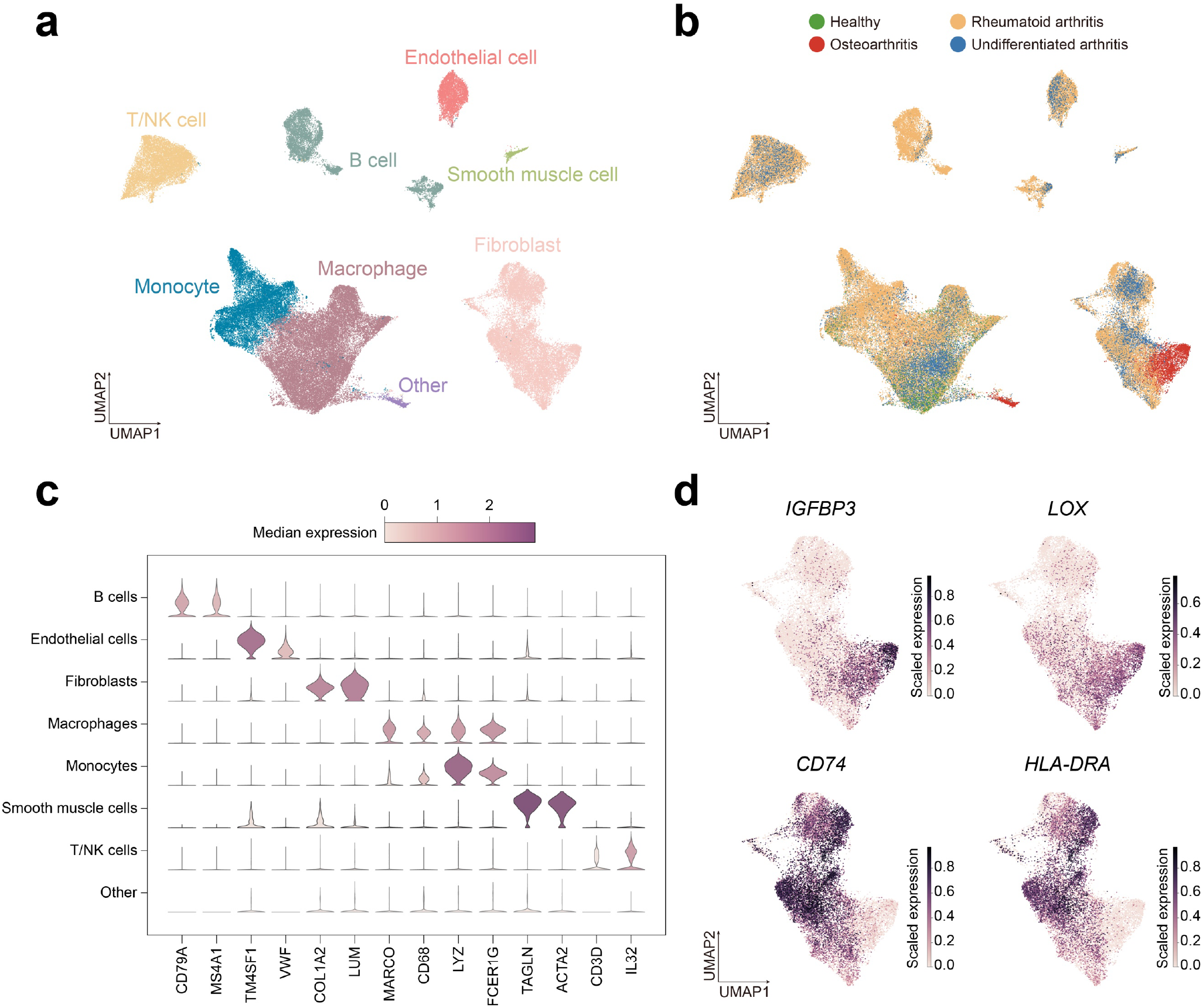
Integrated analysis of scRNA-seq data of synovial tissues-derived cells from OA, RA, UA and healthy patients. **a, b**, UMAP plots showing clustering of 101,610 cells from healthy, osteoarthritis, rheumatoid arthritis and undifferentiated arthritis patients. Cells are colored by cell types (**a**) and by diseases (**b**). **c**, Violin plots showing marker gene expression of each cell type. **d**, Scatter plot showing the expression level of genes specifically expressed in RA-or OA-derived fibroblasts. Cells are colored by the expression of the indicated genes.

Overall, MSdb is a resource created for the MSK research community and aims to fulfill the findability, accessibility, interoperability, and reusability (FAIR) principles of scholarly data^13^. The uniformity of sample information in MSdb enables metadata-based and database-scale analysis. MSdb’s utility will continue to grow as public NGS data sets of human musculoskeletal system expand. We envision that it will broaden the use of human MSK data sets and will be invaluable to researchers in the MSK field.

## Methods

### Data collection and meta information curation

Bulk RNA-seq, microRNAs-seq and single-cell RNA-seq data of human musculoskeletal system were originated from NCBI Gene Expression Omnibus (GEO) and EMBL’s European Bioinformatics Institute (EMBL-EBI)^14,15^. We manually curated both GEO and EMBL-EBI-derived sample information to provide a coherent and standardized metadata. The resulting collection offers the following information for each dataset. ‘SampleName’ contains the sample’s identification code in GEO (e.g. GSM2112324) or EMBL-EBI (e.g. ERS1034560). ‘ProjectID’ contains the sample’s project identification code in GEO (e.g. GSE80072) or EMBL-EBI (e.g. E-MTAB-4304). ‘Publication_DOI’ contains the digital object identifier of the original publication. ‘Category’ indicates the MSK tissues that the datasets are related to. ‘AssayType’ indicates which sequencing types were implemented on the samples. ‘LibraryLayout’ refers to pair-end sequencing or single-end sequencing. ‘Disease’ contains the information about the diseases or health status. ‘SourceTissue_type’ indicates tissue sources of the biological materials used for sequencing. ‘Sourcetissue_condition’ indicates whether the tissues are pathological or normal. ‘SourceTissue_position’ refers to a more specific anatomical location of the ‘SourceTissue_type’. ‘SourceTissue_celltype’ indicates whether whole tissue or a specific cell type in the tissue was used for sequencing. ‘OtherInfo’ contains other information that can help users to evaluate the biological or clinical relevance of the data, such as whether the patients were response to treatment. ‘Age’, ‘AgeGroup’ and ‘Gender’ of the patients are also presented, if available. To assist evaluating the quality of the datasets, we also provide the following information related to data quality assessment along with metadata: sequencing library preparation kit (‘LibraryPrepKit’), average spot length (‘AvgSpotLen), sequencing instrument (‘Instrument’), total reads (‘TotalReads’), uniquely mapped reads (‘UniquelyMappedReads’), percentage of uniquely mapped reads (‘UniquelyMappedReads%’), percentage of multiple mapped loci (‘MultipleLoci%’).

### Bulk RNA-seq and microRNA-seq processing and data analysis

Quality control (QC) of raw sequencing reads for each project was performed by FastQC (v0.11.9)^16^. Cutadapt (v3.7) was used to find and remove adapter sequences, primers, low quality sequence and other types of unmated sequences^17^. For bulk RNA-seq, the trimmed reads were mapped to the reference index built on human genome assembly GRCh38 (hg38, http://ensembl.org/Homo_sapiens/) using STAR (v2.7.9a) and counts were summarized to the genomic protein coding genes by featureCounts (v2.0.3)^18,19^. For microRNA-seq, trimmed reads were mapped to the reference index built on miRbase hairpins and Samtools (v1.14) was used to report alignment summary statistics and calculate the microRNA counts^20^. Batch effect between different studies was estimated and adjusted by ComBat-seq^21^. Then, all bulk RNA-seq gene counts and microRNA-seq counts were merged, respectively. To show sample correlation, Scanpy (1.9.1) was used to reduce dimension and generate uniform manifold approximation and projection (UMAP)^22^. To perform differential expression analysis, a t-test was applied to the normalized RPKM or RPM data and a false discovery rate (FDR) adjusted p-value was canulated using Benjamini–Hochberg method。

The MSdb’s online differential expression analysis tool (https://www.msdb.org.cn/browse/) was used to obtain differentially expressed mRNAs and miRNAs (FDR < 0.01 and fold change > 2). Down-regulated microRNAs and up-regulated genes in degenerative intervertebral disks were used for further analysis. CyTargetLinker (v4.1.0) was used to predict and construct the miRNA-gene interaction network with miRTarBase Homo sapiens release 8.0 linksets was used as a reference^23,24^. Cytoscape software (v3.9.1) was used for miRNA-mRNA regulatory network visualization^25^.

### Single cell RNA-seq processing and data analysis

The genome reference used in scRNA-seq analysis is GRCh38. For droplet-based scRNA-seq, the raw data were processed using Cell Ranger (v7.0.0) or Drop-seq_tools with standard pipeline and default parameters to obtain gene expression matrix^26,27^. For full-length scRNA-seq, the data were mapped using STAR (v2.7.9a) and quantified using featureCounts (v2.0.3). To perform downstream analysis, the gene expression matrix containing UMI counts was read into an AnnData object by Scanpy (v1.9.1) in Python3 (v3.9.12)^22^. Cells with unique gene counts less than 200 or genes that are detected in less than 3 cells were removed. To perform unsupervised cell clustering analysis, the UMI counts were normalized to counts per million (CPM) with *scanpy*.*pp*.*normalize_total* function, followed by log-transformation and principal component analysis (PCA) using *scanpy*.*pp*.*log1p* and *scanpy*.*tl*.*pca* functions. The neighborhood graph was calculated based on the PCA results using *scanpy*.*pp*.*neighbors* function and the Leiden algorithm was used to perform unsupervised cell clustering (*scanpy*.*tl*.*leiden*)^28^. Marker genes for each cell cluster were identified by *sc*.*tl*.*rank_genes_groups* function. To perform reference-based cell subtype prediction, the filtered count matrix was loaded into Seurat (v4.1.1) in R (v4.2.0)^29^. The annotation of cell subtype was performed by SingleR (v1.10.0) R-package using different references, including the BlueprintEncodeData and the HumanPrimaryCellAtlasData^30^. Marker genes for each annotated cell types were identified by *FindAllMarkers* in Seurat package. UMAP was used for the data visualization of unsupervised cell clustering, cell type prediction and marker gene expression.

We built a single-cell atlas of the synovium containing 101,610 cells from 3 studies and 34 samples. We have implemented a probabilistic model based on a variational autoencoder to integrate single-cell RNA-seq datasets and remove batch effects, accepting raw count matrix as input. The variational distribution adopts the log-normal distribution with scalar mean and variance output from the encoder, regularized by the Kullback–Leibler divergence. The decoder takes categorical encoding of the sample name to reflect biological variance and remove batch effects. The count data is modelled by the zero-inflated binomial distribution. The dimension of the latent embedding of the variational autoencoder was chosen to be 10. The top 3,000 highly variable genes were selected using Scanpy (v1.9.1) for the model to learn the latent embeddings^22^. The model was trained on NVIDIA GeForce RTX 3090 addressing 24 GB RAM. Cell annotations for the integrated datasets were based on unsupervised clustering result and prior knowledge of marker gene expression of major cell types including B cells, T/NK cells, macrophages, monocytes, fibroblasts, endothelial cells and smooth muscle cells. The final representation of the dataset was projected to 2-dimensional space using the UMAP algorithm.

### Website development

We developed a user-friendly web interface with advanced functions to present our uniformly curated metadata and NGS data. The front-end interface was developed with HTML5 and CSS3 languages, based on the BootStrap (v5.2.1) toolkit. All font-end tables were built through DataTables (v1.12.1), a plug-in for the jQuery *Javascript* library. All data visualizations were developed by D3.js (v7), a JavaScript library for document manipulation. All back-end data including the bulk RNA-seq and microRNA-seq gene count matrix, UMAP coordinate information, the scRNA-seq clusters, cell subtype annotation, marker genes for different clusters were maintained into PostgreSQL database management system (v14.5). The MSdb database is deployed with a Nginx web server (v1.18.0) on an Ubuntu Linux (v20.04.5 LTS) operating system.

## Acknowledgements

The authors would like to thank all researchers who generated the data sets that are collected, analyzed, and displayed in MSdb. We thank Dr. Xiao Chen, Dr. Zi Yin, Dr. Wenyan Zhou, Dr. Dr. Can Zhang, Dr. Xiaolei Zhang, Fan Jiayi, and clinicians Yan Wu (M.D.), Dr. Yejun Hu (M.D.), Dr. Kun Zhao (M.D.), Dr. Yuzi Xu (M.D.) and Dr. Geyu Gu (M.D.) for their helpful discussions and valuable suggestions. We would also like to thank the technical support provided by the Core Facilities, especially the ZJE server of ZJU-UoE Institute. This work was funded by National Natural Science Foundation of China [32170551 to L.W.; T2121004 to H.O.]; Fundamental Research Funds for the Central Universities 226-2022-00134 (to W.L.); Alibaba Cloud [to L.W.].

## Author contributions

J.L., W.L., H.O., and W.S. conceived the study and designed database. R.T., P.C., C.D., and J.L. collected and processed the data. R.T., D.R., Y.X. and J.L. curated the metadata. R.T. and Z.X. developed the database and web server. R.T., Z.X., D.R., J.L., and W.L. analyzed the data. R.T., Z.X., D.R., W.L. and J.L. wrote the manuscript. All authors contributed to the review and corrections of the manuscripts.

## Data availability

All raw data are available on GEO (https://www.ncbi.nlm.nih.gov/geo/) and EMBL-EBI (https://www.ebi.ac.uk/) repositories. All curated sample information and processed bulk RNA-seq and microRNA-seq data can be downloaded from the MSdb database (https://www.msdb.org.cn). Single-cell RNA-seq matrices are available upon reasonable request.

## Code availability

We used publicly available software to process and analyze data, which are listed and described in Methods. Code for VAE used in this paper for data integration can be found at https://github.com/wanluliu/2022_MSdb_code.

## Competing interests

The authors declare no competing interest.

## Supplementary figures and legends

**Supplementary Fig. 1.**
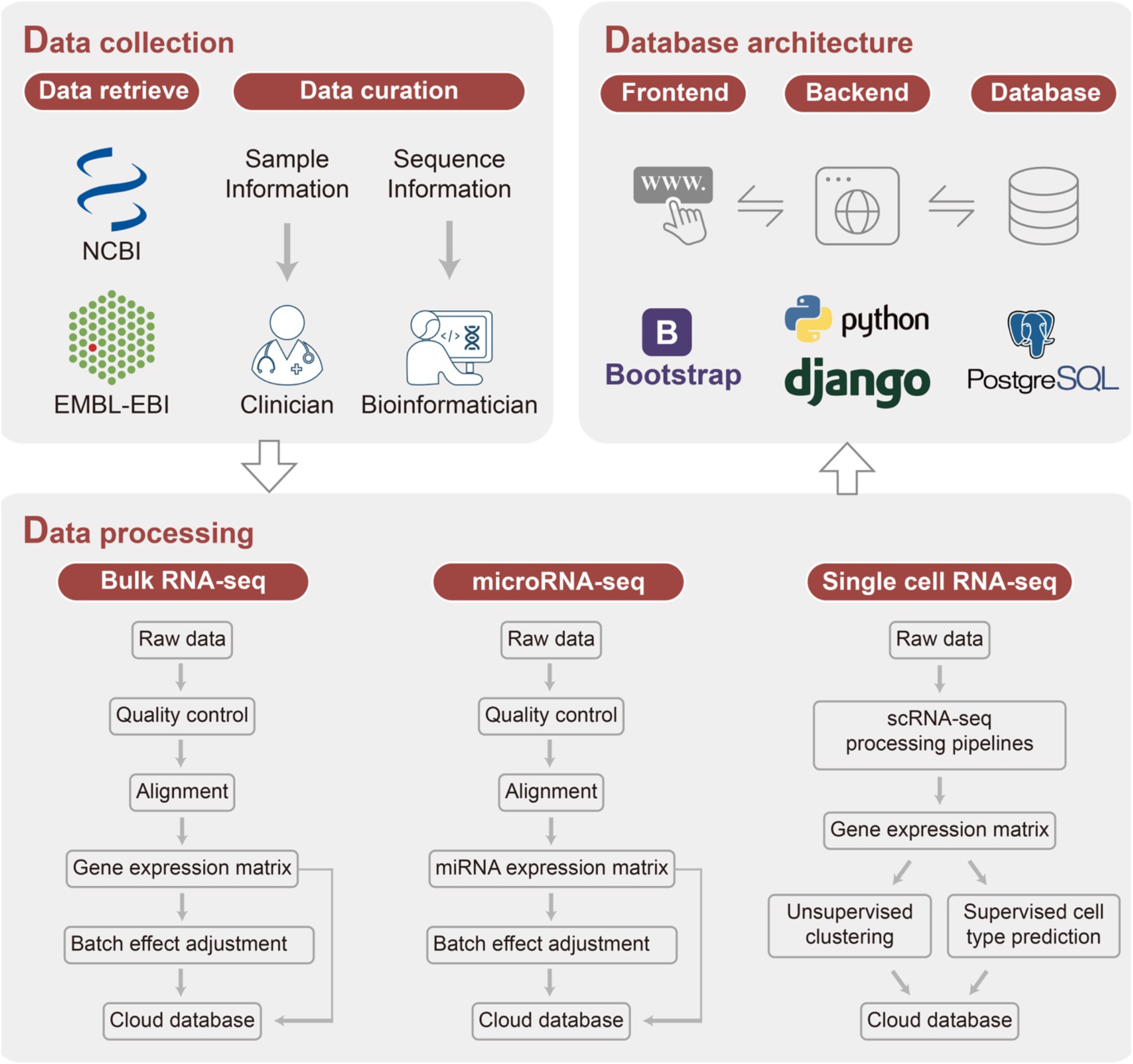
Overview of MSdb database construction. Publicly available data sets were collected from NCBI and EMBL-EBI databases. Metadata was curated by clinician and bioinformatician. Standardized data processing pipelines were built for bulk RNA-seq, microRNA-seq and single-cell RNA-seq. All processed data were stored in PostgreSQL database, and can be accessed with a user-friendly web interface.

**Supplementary Fig. 2.**
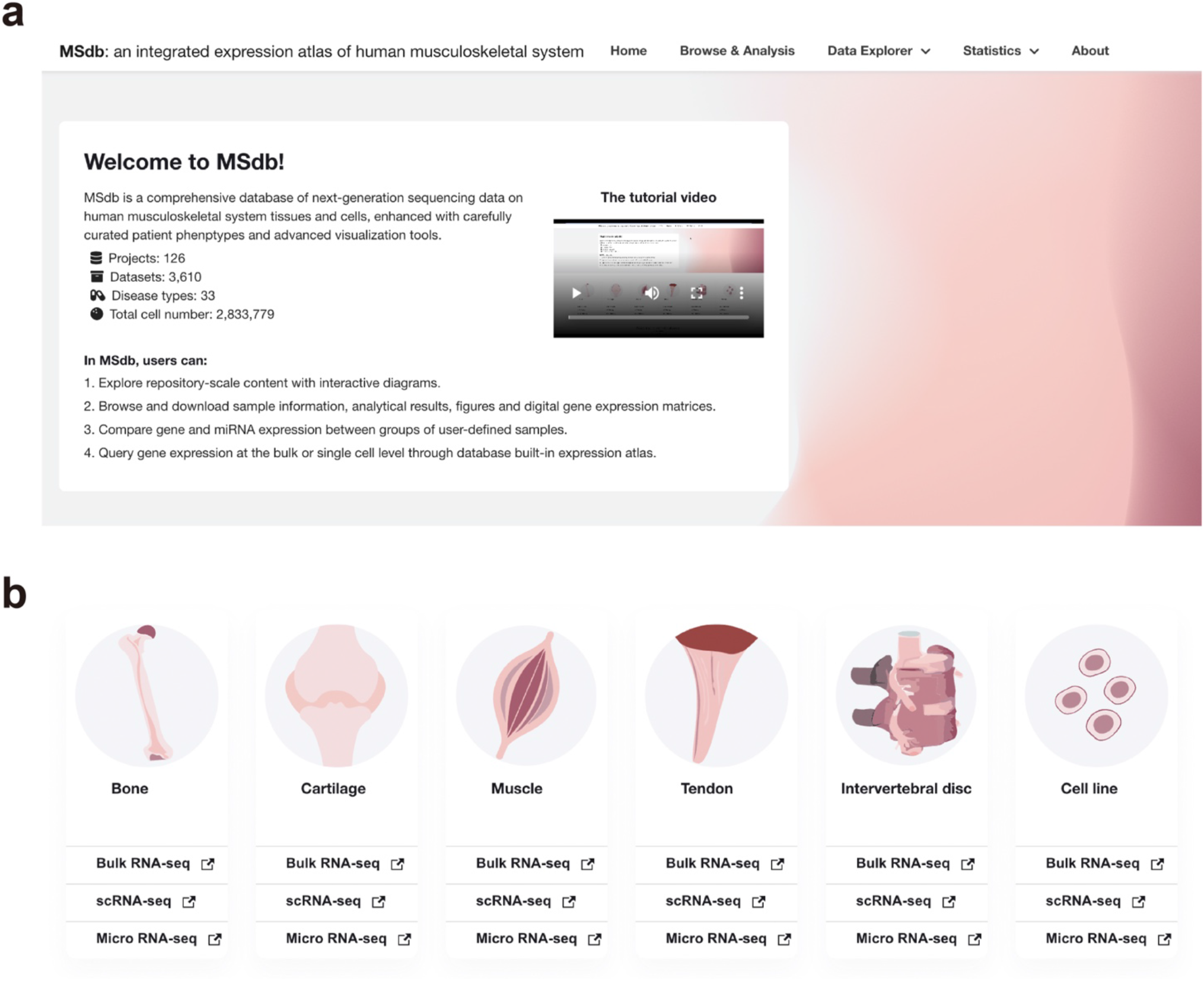
An illustration of the MSdb home page. **a**, The upper portion of the home page showing site-wide menu choices and brief introduction of the database. **b**, The lower portion of the home page showing tissue icons and links for quick access to each data type.

**Supplementary Fig. 3.**
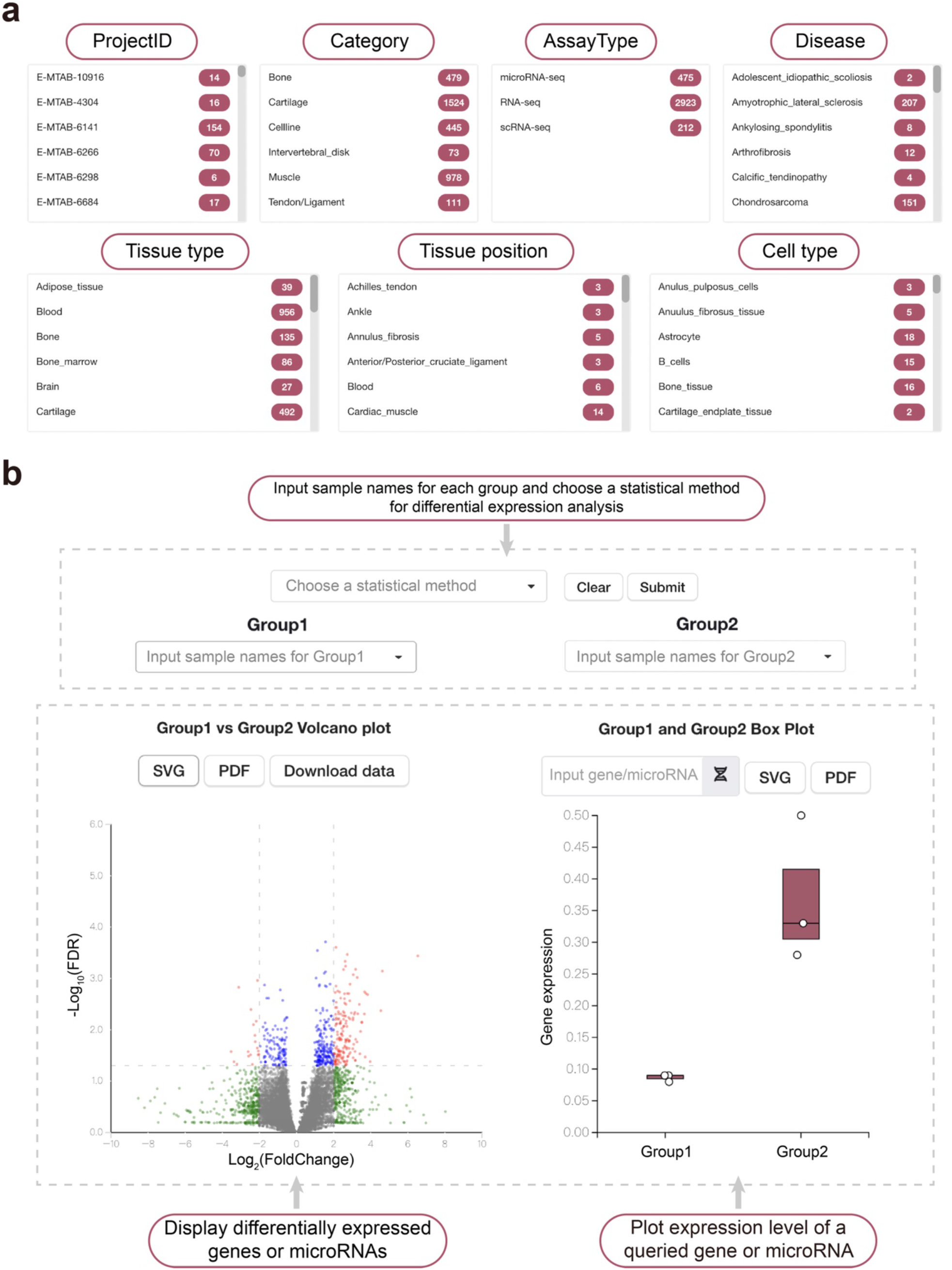
An illustration of browse and differential expression analysis modules. **a**, The selection boxes. Users can select data sets based on the project ID, tissue type, assay type, disease, source tissue type, source tissue position or source tissue cell type. **b**, The differential expression analysis module. Users may choose no more than 8 samples for each group. The differentially expressed genes between the user-defined samples will be calculated and presented in a volcano plot. Users may query the expression level of a specific gene or microRNA in the box plot.

**Supplementary Fig. 4.**
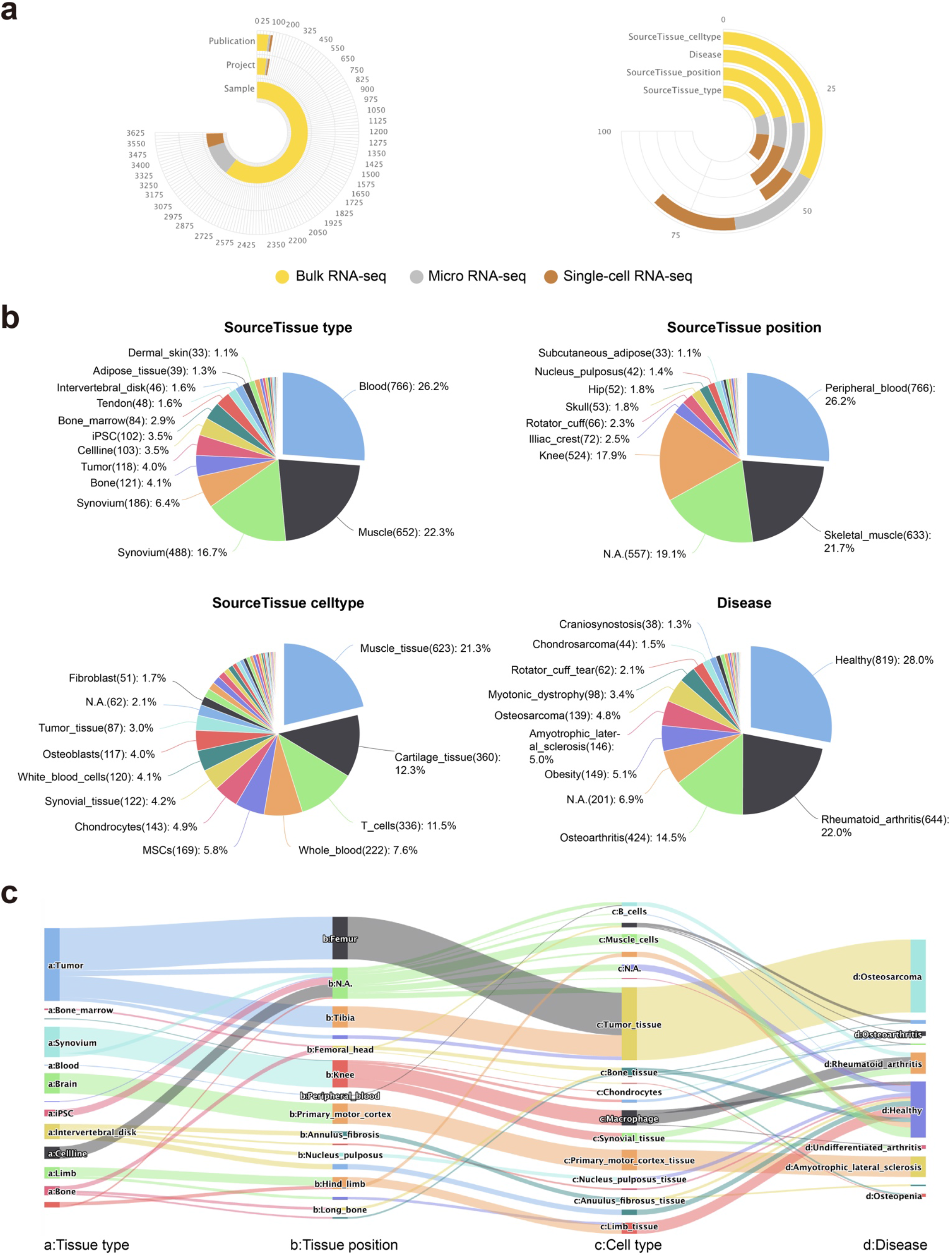
An illustration of data statistics on MSdb. **a**, The circular bar plots showing the number of publications, projects and samples (left), as well as the number of diseases, tissue types, positions and cell types (right) in MSdb. **b**, Pie charts showing the distribution of samples across tissue types, tissue positions, cell types and diseases. Shown are the statistics of bulk RNA-seq, and the statistics of other assay types can be found on our website. **c**, Sankey diagram displaying Tissue type - Tissue position - Cell type - Disease relationships of the samples. Shown is the diagram of scRNA-seq, and the diagrams of other assay types can be found on our website.

**Supplementary Fig. 5.**
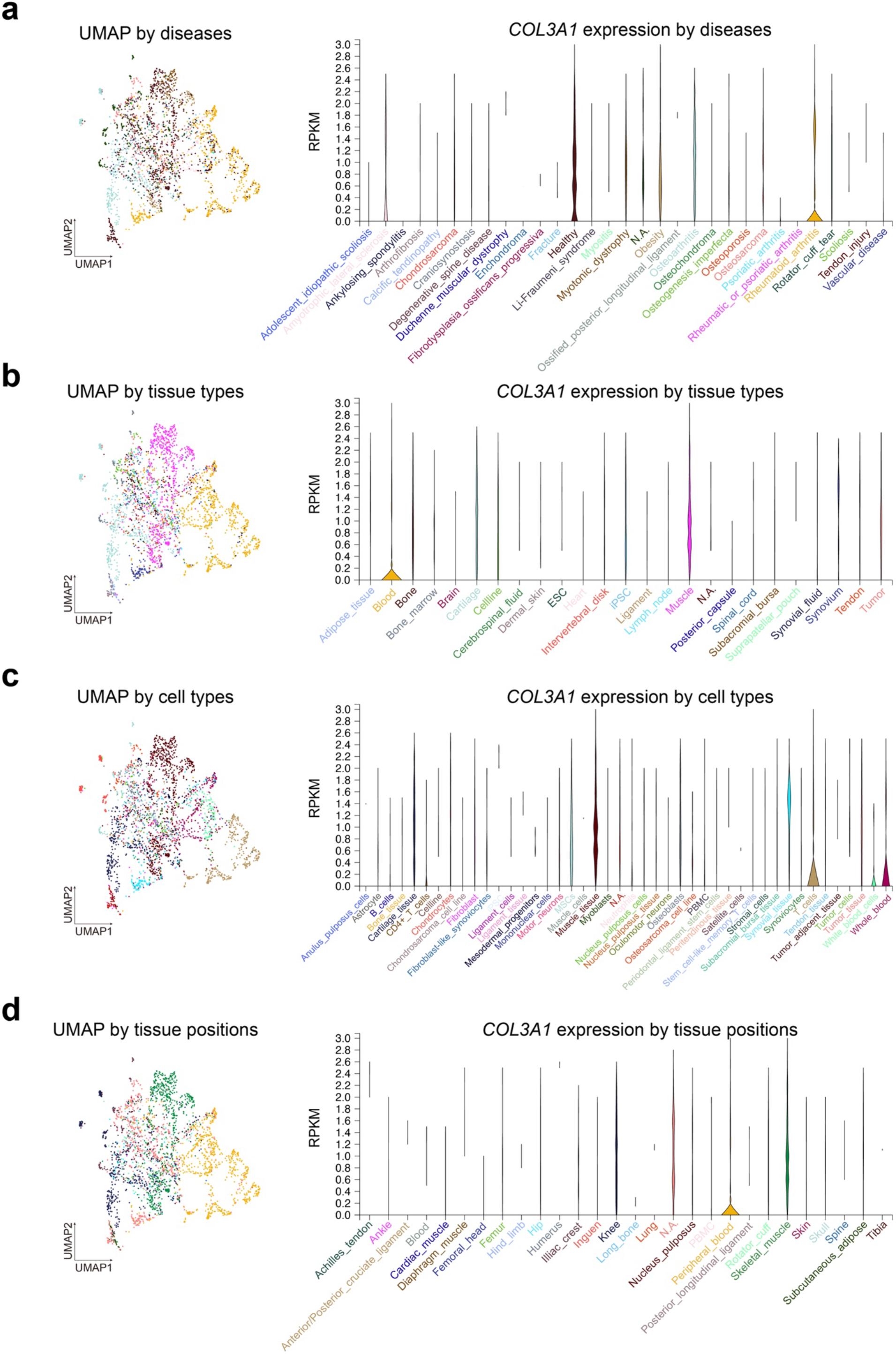
Sample clustering and gene expression plots of bulk RNA-seq. UMAP plots and violin plots showing sample clustering and *COL3A1* expression by diseases (**a**), tissue types (**b**), cell types (**c**) and tissue positions (**d**). The color code of UMAP in each panel are indicated by the colors of the text below the violin plots.

**Supplementary Fig. 6.**
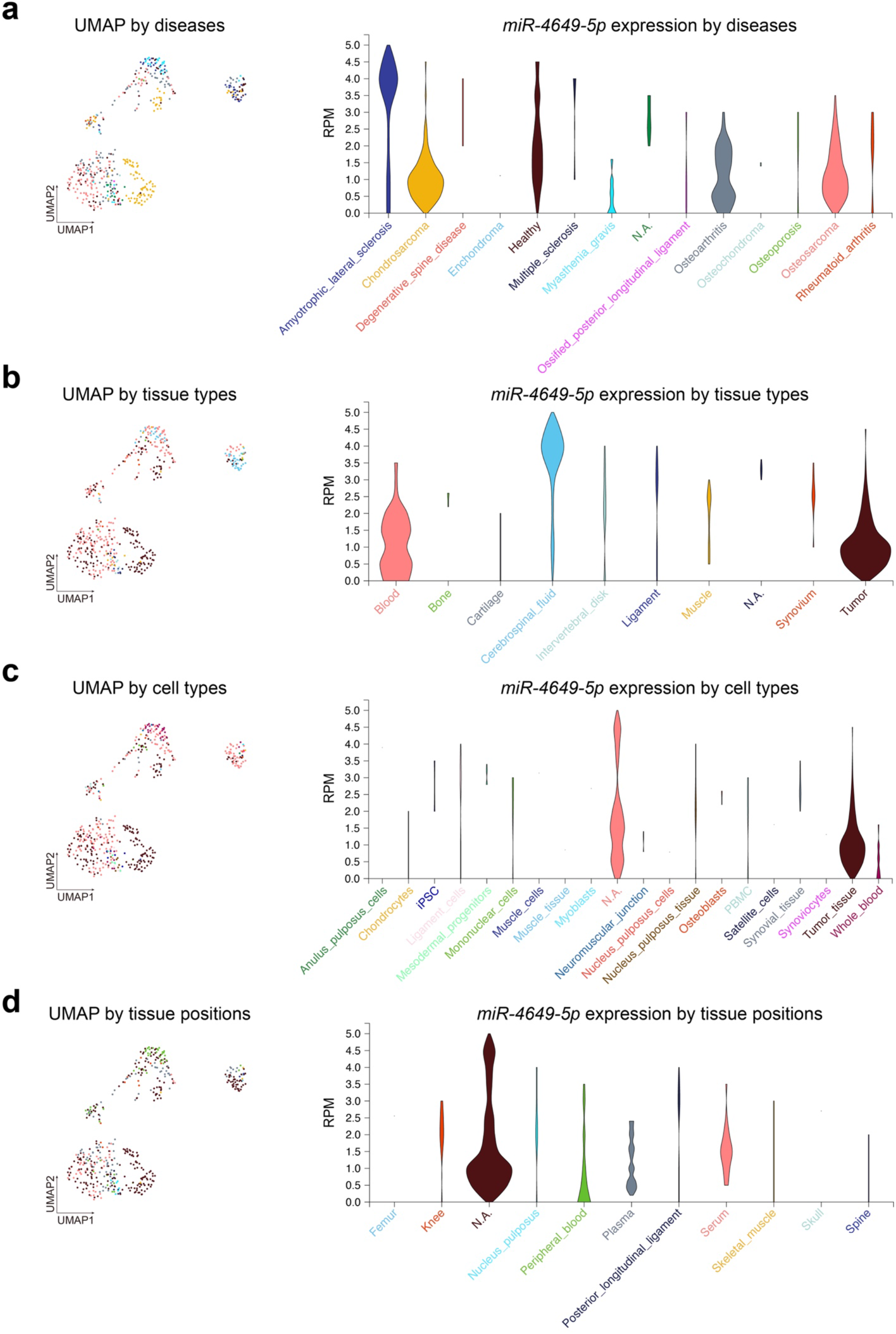
Sample clustering and microRNA expression plots of microRNA-seq. UMAP plots and violin plots showing sample clustering and *miR-4649-5p* expression by diseases (**a**), tissue types (**b**), cell types (**c**) and tissue positions (**d**). The color code of UMAP in each panel are indicated by the colors of the text below the violin plots.

**Supplementary Fig. 7.**
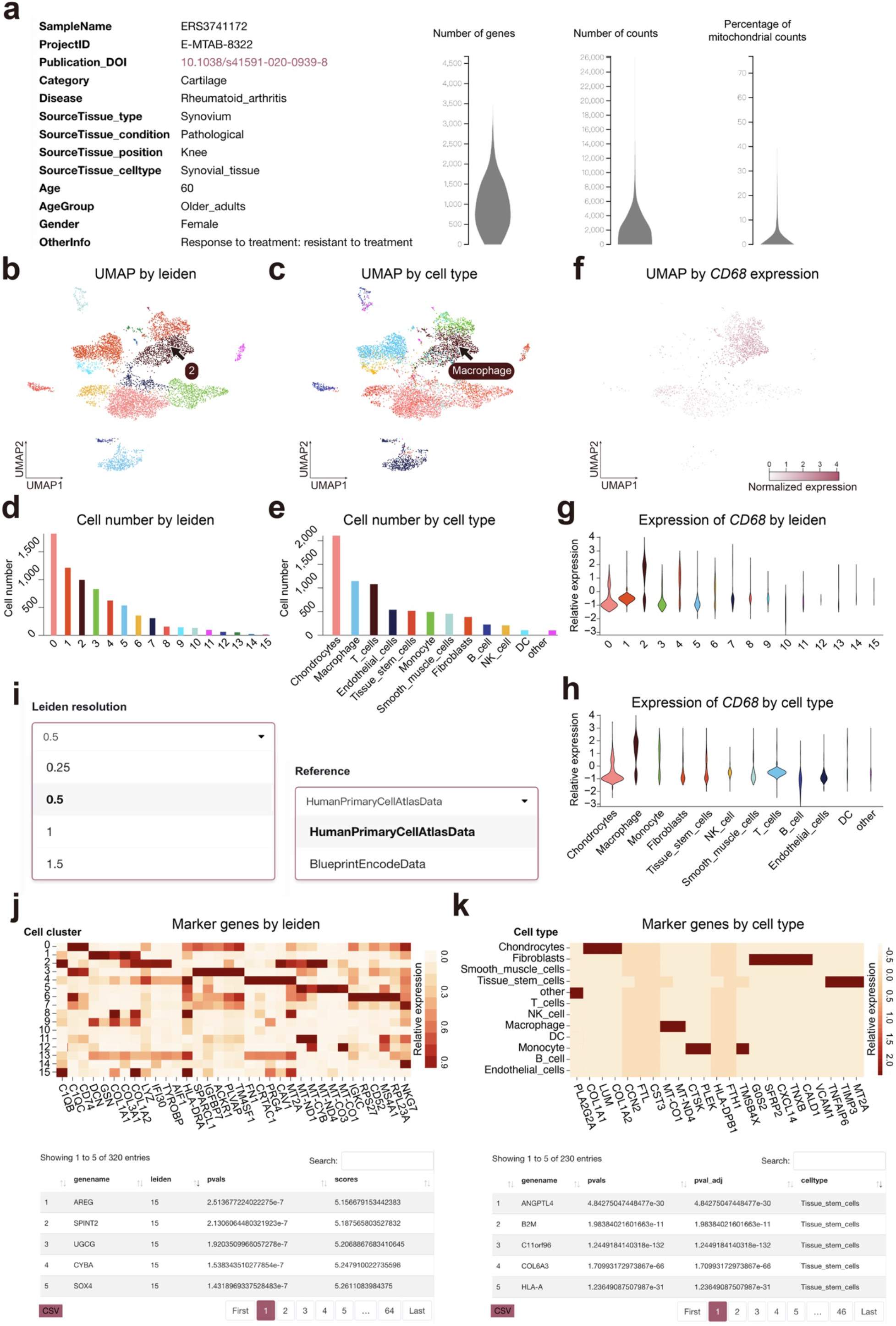
An illustration of scRNA-seq expression atlas in MSdb. **a**, Metadata and quality control metrics of the selected scRNA-seq dataset. **b, c**, UMAP plots showing unsupervised cell clustering (**b**) and cell type prediction (**c**) results. On web interface, the name of the cell cluster or cell type will be shown if one hovers the cursor on the dot. **d, e**, Cell number in each cell cluster (**d**) and predicted cell type (**e**). **f**, UMAP plot showing expression level of *CD68* gene in each single cell. **g, h**, Violin plots showing expression level of *CD68* gene in each cell cluster (**g**) and predicted cell type (**h**). **i**, Dropdown boxes offering choice for Leiden resolution (left) and reference data sets (right) for cell clustering and cell type prediction, respectively. **j, k**, The heatmap and table of marker genes for each cell cluster (**j**) and predicted cell type (**k**). The table can be downloaded in CSV format.

**Supplementary Fig. 8.**
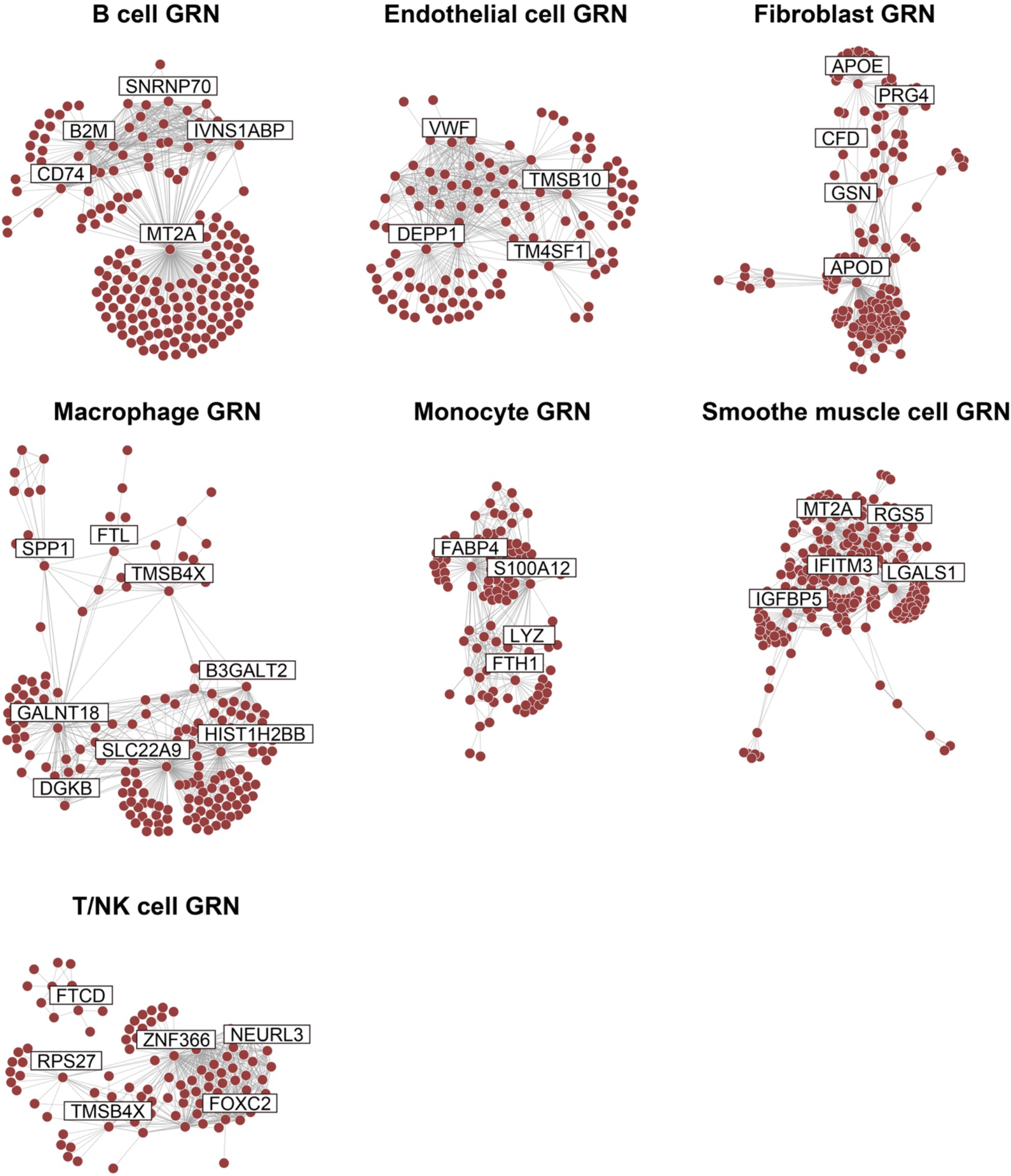
Gene regulatory networks inferred from scRNA-seq. Gene networks for each cell types in Fig. 2a. Red dots represent genes. Core genes are highlighted. Complete and interactive plots of these GRNs are available online on MSdb database.

## Notes

### Competing Interest Statement

The authors have declared no competing interest.

